# REvolutionH-tl: A Fast and Robust Tool for Decoding Evolutionary Gene Histories

**DOI:** 10.1101/2025.04.03.646975

**Authors:** José Antonio Ramírez-Rafael, Annachiara Korchmaros, Katia Aviña-Padilla, Alitzel López-Sánchez, Gabriel Martinez-Medina, Alfredo J. Hernández-Álvarez, Marc Hellmuth, Peter F. Stadler, Maribel Hernández-Rosales

## Abstract

`REvolutionH-tl` is a fast, scalable, and integrated software platform for inferring orthology relationships, gene trees, species trees, and reconciled evolutionary scenarios directly from sequence data. Built upon the formal framework of best match graphs (BMGs), `REvolutionH-tl` predicts orthogroups and orthologous gene pairs with high accuracy, requiring neither precomputed trees nor multiple external tools. The software reconstructs event-labeled gene and species trees, seamlessly integrating reconciliation to produce fast, accurate, and biologically insightful evolutionary scenarios.

Through extensive benchmarking on synthetic datasets with known ground truth, `REvolutionH-tl` outperforms or matches the accuracy of established tools such as OrthoFinder, Proteinortho, RAxML, GeneRax, and RANGER-DTL, while achieving significantly lower runtimes. A key innovation of `REvolutionH-tl` is its built-in support for detailed, publication-ready visualizations, which allow users to explore genome evolution dynamics, orthogroup composition, and reconciliation results with clarity and ease. These visual features position `REvolutionH-tl` as the first platform of its kind to combine analytical precision with intuitive interpretability. The software is open-source, cross-platform, and freely available at https://pypi.org/project/revolutionhtl/, providing a robust solution for large-scale evolutionary analyses in comparative genomics.

**Author summary:** Comparative genomics relies on understanding how genes evolve across species. This involves identifying groups of related genes, reconstructing their evolutionary trees, and aligning them with the evolutionary history of species. These steps are typically performed using multiple tools, often requiring manual integration and technical expertise.

We present `REvolutionH-tl`, an open-source software that automates the entire evolutionary reconstruction process—starting from protein sequences and producing gene trees, species trees, orthology assignments, and reconciled evolutionary scenarios. Unlike existing tools, `REvolutionH-tl` also includes built-in, high-quality visualizations that help users interpret complex evolutionary events such as gene duplications and losses.

We evaluated `REvolutionH-tl` on simulated genomes with known evolutionary histories and found that it matches or exceeds the performance of widely used tools, while being significantly faster. Its visual output makes evolutionary analysis more accessible and interpretable, offering a valuable resource for researchers studying genome evolution.

## Introduction

Phylogenetics reconstructs evolutionary histories by tracing shared characteristics in descendant lineages back to a common ancestor. The best-supported hypotheses of evolutionary histories are usually represented with phylogenetic trees, where each node depicts the emergence of a new lineage, while each branch represents the preservation of a genetic lineage over time. In particular, if a tree represents the evolutionary relationship among a group of species, then the branching points represent a speciation event. On the other hand, if a tree represents the evolution of genes, then the internal nodes represent speciation, duplication, or other events the genes went through.

Reconstructing the evolutionary history between genomic entities helps to understand the evolution of morphological characteristics, reconstruct demographic changes in recently diverged species, and infer, in particular, the function of newly discovered genes. Functional gene annotation relies on identifying which homologous genes are orthologs, as they tend to preserve functions that were present in their ancestors [1]. In this context, a *gene* represents either a nucleotide or amino acid sequence, and a set of homologous genes share a common ancestor. Two genes are *orthologs* if they diverge from a single ancestral gene amid the speciation process of the last common ancestor of the species in which such genes reside; orthologs originating from a gene of the last common ancestor of the species under consideration constitute an *orthologous group* (OG). Although genetic material may be transferred between species, giving rise to xenologous genes, in this contribution we assume the evolutionary histories are free of horizontal gene transfers (HGT), i.e., the nodes in the gene tree represent either duplication or speciation events. We refer the reader to [2] for further details on gene transfers.

With increasing attention regarding genome and transcriptome annotation, the development of computational methods to infer orthologous relations and reconstruct the gene and corresponding species trees have gained considerable interest [3, 4]; despite the increasing availability of phylogenetic data, the process of accurately reconstructing evolutionary histories is not always straightforward. Methods for orthology detection are either tree-based or graph-based. Tree-based methods identify orthologs by reconciling an explicit tree model of the genes’ history to one of the species where they reside. Orthologs are then set to be genes that group with members from other species in the tree reconciliation, as in [3]; in [5] this approach is formalized in an algorithm that assigns gene duplication and speciation events in polynomial time. In principle, tree-based methods are the most appropriate for orthologous detection; however, such methods are precluded when the number of leaves is large, as trees are computationally expensive to produce [3].

Our main interest is in graph-based orthology methods. Such methods represent homologous genes as the nodes of a graph whose edges are their orthologous relations estimated with sequence similarity. In HGTs-free scenarios, orthologs are *best matches*, i.e., the homologs from different species that branched last; the vice-versa is not necessarily true if orthologs are lost during speciation and only one gene family member remains in each species [6]. Moreover, if evolutionary closeness is estimated by means of sequence similarity, orthologs are detected as *best hits*, i.e. the genes with the highest similarity score match. The maximum likelihood estimation of evolutionary distances has also been used to detect orthologs [7]. Pairwise sequence similarity scores are usually computed as bit-scores using all-vs-all `BLAST` search [8] or `DIAMOND` [9] as an accelerated alternative to `BLAST`. The bidirectional best hits is the most popular graph-based approach, assuming the orthologous inference is as symmetric as its evolutionary relation counterpart. This approach limits one ortholog per gene and species, which is false when a gene undergoes duplication after speciation. Methods that allow multiple orthologs (co-orthologs) are available; in particular, `ProteinOrtho` [10] selects the orthologs whose score is above a dynamical score that depends on the gene and species considered. Graph-based approaches are computationally efficient and scale well with large datasets compared to tree-based approaches. However, as in tree-based approaches, orthologs paired with reconciled gene trees provide more information to the user than sets of orthologs. An alternative to the classical methods for assessing orthology in sequenced transcriptomes is comparing newly discovered sequences to a set of reference orthologs from dedicated databases [11].

Phylogeny reconstruction methods are either distance-based or character-based. Distance-based methods iteratively build a tree using a distance matrix between every pair of taxa. Neighbour joining [12] is the most popular of such methods, and recent implementations typically run approximately to the square of the number of taxa; however, the correctness of the output relies on an additive and unbiased distance matrix, which cannot also be guaranteed. Maximum parsimony (MP), maximum likelihood (ML), and Bayesian inference (BI) are the main types of character-based methods [13]. BI and ML rely on an explicit evolutionary model and the likelihood function *L*(*θ*), which is the probability of observing the data given that the parameter *θ* includes substitution model and branch length parameters. The ML tree estimates the topology with the highest likelihood. BI tree estimate is instead the maximum posterior probability tree among all topologies simulated via the Markov chain Monte Carlo algorithm, where the posterior probability is obtained by multiplying a prior distribution and the likelihood. To avoid parameter estimations of a given evolutionary model, recently, phylogenetic invariants have been used to estimate the tree topology [14, 15]. All these methods are computationally expensive and require specific evolutionary model assumptions, which may negatively affect the tree estimate. On the other hand, the maximum parsimony (MP) approach, which is the one of interest in this paper, is not susceptible to the drawbacks of likelihood-based models; indeed, MP tree estimation involves the minimum number of events among the taxa. However, MP is more susceptible to the long-branch attraction artifact (LBA) in the case of heterogeneous rates.

Susceptibility to LBA may be caused by the theoretical gap between best hits and best matches; indeed, pairs of taxa with the highest bit-scores are not necessarily the closest evolutionary as in [4]. This observation motivated a recent mathematical interest [16–20] in detecting orthology through best match graphs, which group the most closely related genes between species in directed graphs called BMGs. In [21], a theoretical pipeline for inferring orthologous relationships and tree phylogenies was proposed but remained unimplemented until recently. Ramírez-Rafael et al. [22] introduced `REvolutionH-tl` (**R**econstruction of **Evolution**ary **H**istories **t**oo**l**), a novel bioinformatics tool that applies the BMG theoretical framework to identify orthologous genes and groups (orthogroups) and to reconstruct and reconcile event-labeled gene and species trees in the presence of duplications and losses.

`REvolutionH-tl` belongs to the family of graph-based orthology methods, initially constructing a BMG estimate from best-hit data (e.g., `BLAST`, `Diamond`). Gene and species trees are inferred by maximizing the number of informative triples—BMG building blocks—without relying on explicit evolutionary models, aligning the method with maximum parsimony principles. Reconciliation is performed by mapping duplications and losses onto species tree branches, while contradictions between gene and species trees prompt alternative hypotheses through gene tree editing, enabling the detection of pseudo-orthologs and pseudo-paralogs.

Here, we present a new version of `REvolutionH-tl`, an enhanced platform of the original tool that incorporates improved components and data structures, resulting in significantly faster performance. In addition, it introduces a modified neighbor-joining method to resolve duplication nodes in gene trees. This new version also includes a comprehensive set of visualizations designed to facilitate more user-friendly analysis and interpretation of results.

We evaluated `REvolutionH-tl` against established tools such as `ProteinOrtho` [10, 23, 24], `OrthoFinder` [25, 26], `GeneRax` [27], `ASTRAL-Pro` [28], `RAxML` [29], and `RANGER-DTL` [30]. Our benchmarking demonstrates that most of the time `REvolutionH-tl` outperforms in orthology inference, gene and species tree reconstruction while significantly reducing computational time. Its combination of accuracy, efficiency, and scalability makes `REvolutionH-tl` a powerful tool for evolutionary analyses, helping researchers uncover the complexities of gene and species evolution with precision and ease.

## Theoretical Background

### Graph theory preliminaries

A **graph** *G* is an ordered pair *G* = (*V, E*), where *V* is the set of **nodes** and *E* `⊆`{*uv* | *u, v V, u* ≠ *v*} is the set of **edges**, representing connections between nodes. If *uv* `∈` *E* implies *vu* `∈` *E* for all *u, v* `∈` *V*, we call *G* **undirected** and, otherwise, **directed**.

The **degree** of a node *v* `∈` *V* in an undirected graph is the number of edges incident to *v*. In a digraph, the **in-degree** and **out-degree** of *v* refer to the number of incoming edges *uv* and outgoing edges *vu*, denoted 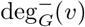 and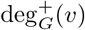, respectively. A **subgraph** *H* of *G* is a graph such that *V* (*H*) `⊆`*V* (*G*) and *E*(*H*) `⊆`*E*(*G*). The **induced subgraph** of *G* on a node set *V* `′ ⊆`*V* (*G*), denoted *G*[*V* `′`], is the subgraph *H* = (*V* `′`, *E*`′`), where *E*`′` = {*uv* `∈` *E*(*G*) | *u, v* `∈` *V* `′`} contains all those edges from *E*(*G*) that connect pairs of nodes in *V* `′`.

A **tree**, *T* = (*V, E*), is a connected and acyclic graph, i.e., removing any edge from *T* disconnects it. A **rooted tree** *T* is a directed acyclic graph (DAG) that does not contain nodes *v* with deg 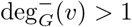. In particular, we assume that *T* contains a unique node *ρ*_*T*_, called **root**, that has no incoming edges and from which all nodes *v* are reachable via a directed path from *ρ*_*T*_ to *v*. By definition, if |*V* | *>* 1, then for each *v* `∈` *V* \ {*ρ*_*T*_} there is a unique incoming edge *uv* and we call *u* the **parent** *p*(*v*) of *v* and *v* a **child** of *u*. We collect in ch_*T*_ (*u*) all childen of *u* in *T*. The nodes of a rooted tree can be classified as **leaves**, *L*(*T*), which are terminal nodes (nodes with no children), and **internal nodes**, *V* ^0^(*T*), which have at least one child. If there is a directed path from *u* to *v*, then *u* is an **ancestor** of *v* and *v* a **descendant** of *u*, denoted by *v* `⪯`_*T*_ *u*. If *v* `⪯`_*T*_ *u* and *u*≠*v*, then we write *v* `≺`_*T*_ *u*. We say that nodes *u* and *v* are **comparable** if *v* `⪯`_*T*_ *u* or *u* `⪯`_*T*_ *v* and, otherwise, we say that *u* and *v* **incomparable** and denote this by *u* `∥` *v*. Furthermore, in [31, 32] the `⪯`_*T*_ relationship has been extended to consider edges within *T* : for edges *e*_0_ = *uv, e*_1_ = *xy* and a node *z, e*_0_ `⪯`_*T*_ *e*_1_ if *v* `⪯`_*T*_ *y, z* `≺`_*T*_ *e*_0_ if *z* `⪯`_*T*_ *v*, and *e*_0_ `≺`_*T*_ *z* if *u* `⪯`_*T*_ *z*.

For any node *v* `∈` *V* (*T*), the **subtree rooted at** *v*, denoted *T* (*v*), is the subgraph of *T* induced by *v* and all its descendants. For any node *v* `∈` *V* (*T*) in a rooted tree *T*, the **cluster** of *v*, denoted *C*(*v*), is the set of leaves in the subtree rooted at *v*, i.e., *C*(*v*) = *L*(*T* (*v*)). The **last common ancestor** (LCA) of a set *X* `⊆` *V* (*T*), denoted lca_*T*_ (*X*), is the node in *T* that is an ancestor of all nodes in *X* and none of the nodes *w* `≺`_*T*_ lca_*T*_ (*X*) satisfy this property. If *X* = {*x, y*} we write lca_*T*_ (*x, y*) instead of lca_*T*_ ({*x, y*}).

A **phylogenetic tree** is a rooted tree where each internal node has at least two children. A **planted tree** is formed by adding a new node 0_*T*_ to a phylogenetic tree *T* and an edge 0_*T*_ *ρ*_*T*_. In a planted tree, the node 0_*T*_ has degree one, with its only neighbor being *ρ*_*T*_, and this property remains unchanged during any subsequent modifications, such as resolving ‘polytomies’. A **polytomy** in a phylogenetic tree is a node with more than two children. Resolving a polytomy involves converting it into a binary tree, a tree where each internal node has exactly two children. This may introduce additional internal nodes to maintain the tree’s structure.

A **triple** *xy*|*z* is a phylogenetic tree *T* on three leaves *x, y*, and *z* such that lca_*T*_ (*x, y*) `≺`_*T*_ lca_*T*_ ({*x, y, z*}). Similarly, a rooted tree *T* **displays** a triple *xy*|*z* if lca_*T*_ (*x, y*) `≺`_*T*_ lca_*T*_ ({*x, y, z*}). Such triples are crucial for reconstructing phylogenetic trees; see, e.g., [33–35]. For any rooted tree *T*, let *R*(*T*) denote the collection of triples displayed by *T*. An arbitrary set *R* of triples is **compatible** if there exists a rooted tree *T* such that *R* `⊆`*R*(*T*). A polynomial-time procedure to verify the compatibility of a set *R* and, if compatible, to construct a tree that displays all triples in *R* is provided by the `BUILD` algorithm [33, 36]. This algorithm constructs an auxiliary graph *G*_*R*,*L*_, commonly referred to as the **Aho graph**, where the nodes correspond to all leaves in *L*, and edges *xy* exist for each triple *xy*|*z* `∈` *R* with *x, y, z* `∈` *L*. This algorithm begins with the set *L* of all leaves in *R* and recurses on the connected components *L*_1_, …, *L*_*k*_ of the Aho graph, repeating the process for the graphs 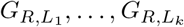. If, at any step, one of the Aho graphs *G*_*R*,*L*_*′* is connected and contains more than one node, the initial set *R* is declared incompatible. Let aho(*R*) be a function that takes as input a triples set *R*, and outputs either the three *T* constructed by `BUILD` or the sentence *the set of triples R is not compatible*.

### Evolutionary scenarios

In what follows we consider particular types of phylogenetic trees, namely species trees and gene trees. A **species tree** *S* = (*V, E*) represents the evolutionary history of a set of species, where the leaves *L*(*S*) `⊆`*V* (*S*) correspond to extant species. A **gene tree** *T* = (*V, E*) represents the evolutionary history of a set of genes, where the leaves *L*(*T*) `⊆`*V* (*T*) correspond to the sampled genes.

Genes evolve within genomes, i.e., species. We denote by *σ* : *L*(*T*) `→` *L*(*S*) the map that assigns each leaf in the gene tree to the species in which it resides. Moreover, we can assign evolutionary events or mechanisms to the nodes of a gene tree *T* that act on genes through evolution. Specifically, the map *t* : *V* (*T*), `→ {`•,□,`⊙`,×} classifies nodes in the gene tree based on evolutionary events: *t*(*x*) = • for speciation, *t*(*x*) = □ for duplication, *t*(*x*) = `⊙` for extant genes, and *t*(*x*) = × for gene loss (see [31, 32], for further details see Fig. 1). Two distinct genes *x, y* `∈` *L*(*T*) are **orthologs** if *t*(lca_*T*_ (*x, y*)) = •. Conversely, they are **paralogs** if *t*(lca_*T*_ (*x, y*)) = □. The inference of such evolutionary events can be approached in two ways. On the one hand, methods based on reconciling the gene tree with a known species tree are used [26, 27, 30, 37]. On the other hand, tree-free methods enable the inference of orthologous and paralogous gene pairs without requiring knowledge of the gene tree or species tree, see [38] for an overview.

**Fig 1.**
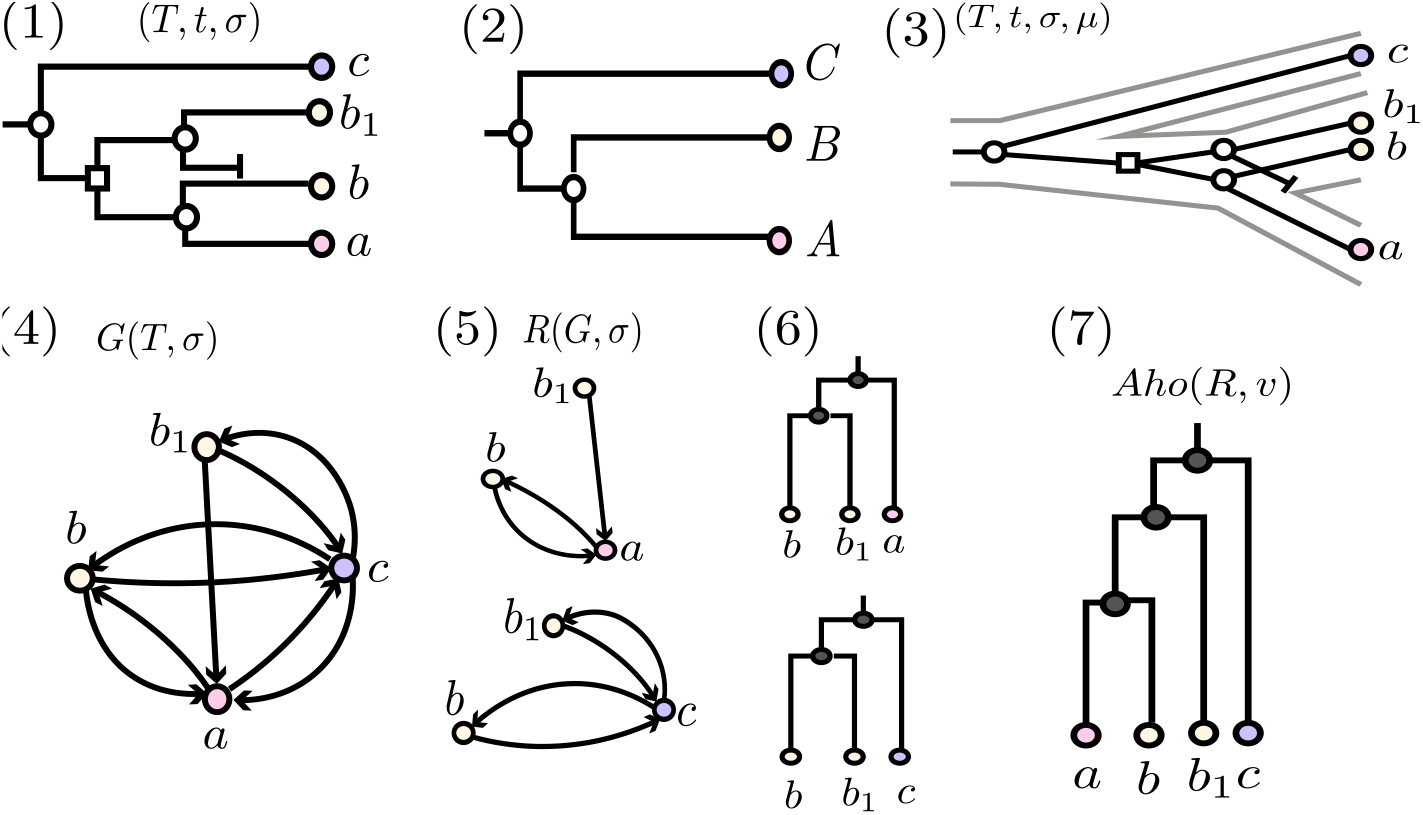
Representation of evolutionary histories and gene tree reconstruction. (1) Event-labeled gene tree (*T, t, σ*) with speciation nodes shown in circles and duplication nodes in squares. Gene-to-species assignments are given by *σ*(*a*) = *A, σ*(*b*) = *σ*(*b*_1_) = *B*, and *σ*(*c*) = *C*. (2) Species tree *S*. (3) Reconciliation map *µ* between the gene tree (*T, t, σ*) and the species tree *S*, illustrated by positioning gene tree nodes within corresponding nodes or edges of *S*. (4) Best match graph *G*(*T, σ*) constructed from the gene tree. (5) Informative triples *R*(*G, σ*) derived from the best match graph. Aho graph based on the triples in *R*(*G, σ*). (7) Tree reconstructed by the `BUILD` algorithm from the set of informative triples.

An **evolutionary scenario** (*S, T, t, σ, µ*) consists of a species tree *S*, a labeled gene tree (*T, t, σ*) and a **reconciliation map** between *S* and *T*, that is, a map *µ* : *V* (*T*) `→` *V* (*S*) `∪` *E*(*S*) that satisfies:

(U1) *Gene Constrain*. If *t*(*x*) = `⊙`, then *µ*(*x*) = *σ*(*x*) `∈` *L*(*S*)

(U2) *Speciation Constrain*. If *t*(*x*) = •, then *µ*(*x*) = lca_*S*_(*σ*(*C*_*T*_ (*x*))) `∈` *V*^0^(*S*), and *µ*(*y*_0_) `∥`_*S*_ *µ*(*y*_1_) for any two distinct children *y*_0_, *y*_1_ `∈` ch_*T*_ (*x*)

(U3) *Duplication Constrain*. If *t*(*x*) = □, then *µ*(*x*) = *e* `∈` *E*(*S*) and lca_*S*_(*σ*(*C*_*T*_ (*x*))) `≺` *e*

(U4) *Ancestor Constrain*. If *x* `≺`_*T*_ *y*, then *µ*(*x*) `⪯`_*S*_ *µ*(*y*) as defined in [31].

The existence of a reconciliation map between a labeled gene tree (*T, t, σ*) and a species tree is characterized by the compatibility of particular triples displayed in *T*. To be more precise, let ℜ (*T*) = {*r* `∈` *R*(*T*) | *t*(lca_*T*_ (*L*(*r*))) = • and |*σ*(*L*(*r*))| = 3} be the set of all triples *ab*|*c* that are rooted at a speciation event, where the species *σ*(*a*), *σ*(*b*), and *σ*(*c*) are pairwise distinct. Given a triple *ab*|*c* `∈` ℜ (*T*), the corresponding *color triple* is *σ*(*ab*|*c*) = *σ*(*a*)*σ*(*b*)|*σ*(*c*). Finally, let ℜ{_*σ*_(*T*) = *σ*(*r*) | *r* ∈ ℜ (*T*)} be the set of *color triples* of the gene tree.

#### Theorem 1

([31, 32]). *For a given labeled gene tree* (*T, t, σ*), *there exists an evolutionary scenario* (*S, T, t, σ, µ*) *if and only if* ℜ_*σ*_(*T*) *is compatible. In this case, every species tree S that displays all triples in* ℜ_*σ*_(*T*) *can be reconciled with* (*T, t, σ*).

In our framework, neither *T*, *t* nor *S* are given and our aim is to infer evolutionary scenarios for a given set of extant genes that solely relies on best hits, best matches and related concepts that are made more precise below.

### Best match graphs and gene trees

In what follows, we are interested in genes *y* that are the evolutionary “most closely” related ones to a given gene *x*. This concept is formalized through the notion of best matches in a given gene tree (*T, σ*). A gene *y* is a **best match** for a gene *x* in *T* if *x* and *y* reside in distinct species and lca_*T*_ (*x, y*) `⪯` lca_*T*_ (*x, y*`′`) for all genes *y*`′` in the species *σ*(*y*). Given a gene tree (*T, σ*), we define the **best match graph** *G*(*T, σ*) = (*V, E*) as a directed graph with node set *V* = *L*(*T*) and a directed edge *xy* `∈` *E* if *y* is a best match for *x* in (*T, σ*). Given an arbitrary colored digraph *G* and a tree *T* with *L*(*T*) = *V* (*G*), we say *T* **explains** *G* if *G* = *G*(*T, σ*).

The structure of BMGs has been extensively studied in recent years [17, 19, 39, 40]. In particular, as shown by Schaller et al. [41], best matches and their “symmetrized” versions paved the way to infer orthologs decreasing false-positive assignments, provided that the evolutionary history is not distorted by so-called horizontal gene transfer.

A key property of best match graphs is that they encode information about gene tree topology in the so called informative triples; for any directed graph *G* and a node-coloring map *σ* : *V* (*G*) `→` *M*, we say that ℛ(*G, σ*) is the set of **informative triples** *r* = *xy*|*y*`′` with *σ*(*x*) ≠ *σ*(*y*) = *σ*(*y*`′`) such that *xy* `∈` *E*(*G*), *xy*`′` ∉ *E*(*G*). Furthermore, if the gene tree is assumed to be binary, ℛ(*G, σ*) also includes those triples *yy*`′`|*x* whenever *xy, xy*`′ ∈` *E*(*G*). The theorem below provides a theoretical strategy to go from a data-inferred BMG, to a gene tree explaining the graph.

#### Theorem 2

([40]). *A colored digraph* (*G, σ*) *is a BMG if and only if the set of triples* ℛ(*G, σ*) *is compatible and G is explained by T* = *aho*(ℛ (*G, σ*)), *i*.*e*. (*G, σ*) = *G*(*T, σ*).

This result allows us to first, identify if a colored digraph *G* is a BMG. In the affirmative case, we can reconstruct a gene tree by running the `BUILD` algorithm with the informative triples ℛ(*G, σ*), and in the negative case, the Aho-graph is helpful to define an heuristic for editing *G* into a BMG with minimal editions, as proposed in [19].

### REvolutionH-tl**Methodology**

The workflow of `REvolutionH-tl`is divided into 7 steps: (1) *Computation of Alignment Hits*, (2) *BMG Estimation and Orthogroup Detection*, (3) *Gene Tree Reconstruction and Orthology Assignment*, (4) *Polytomy Resolution for Gene Trees*, (5) *Species Tree Reconstruction*, (6) *Tree Reconciliation* and (7) *Visualization of Reconciliation Results*. This methodology is illustrated in Fig. 2.

**Fig 2.**
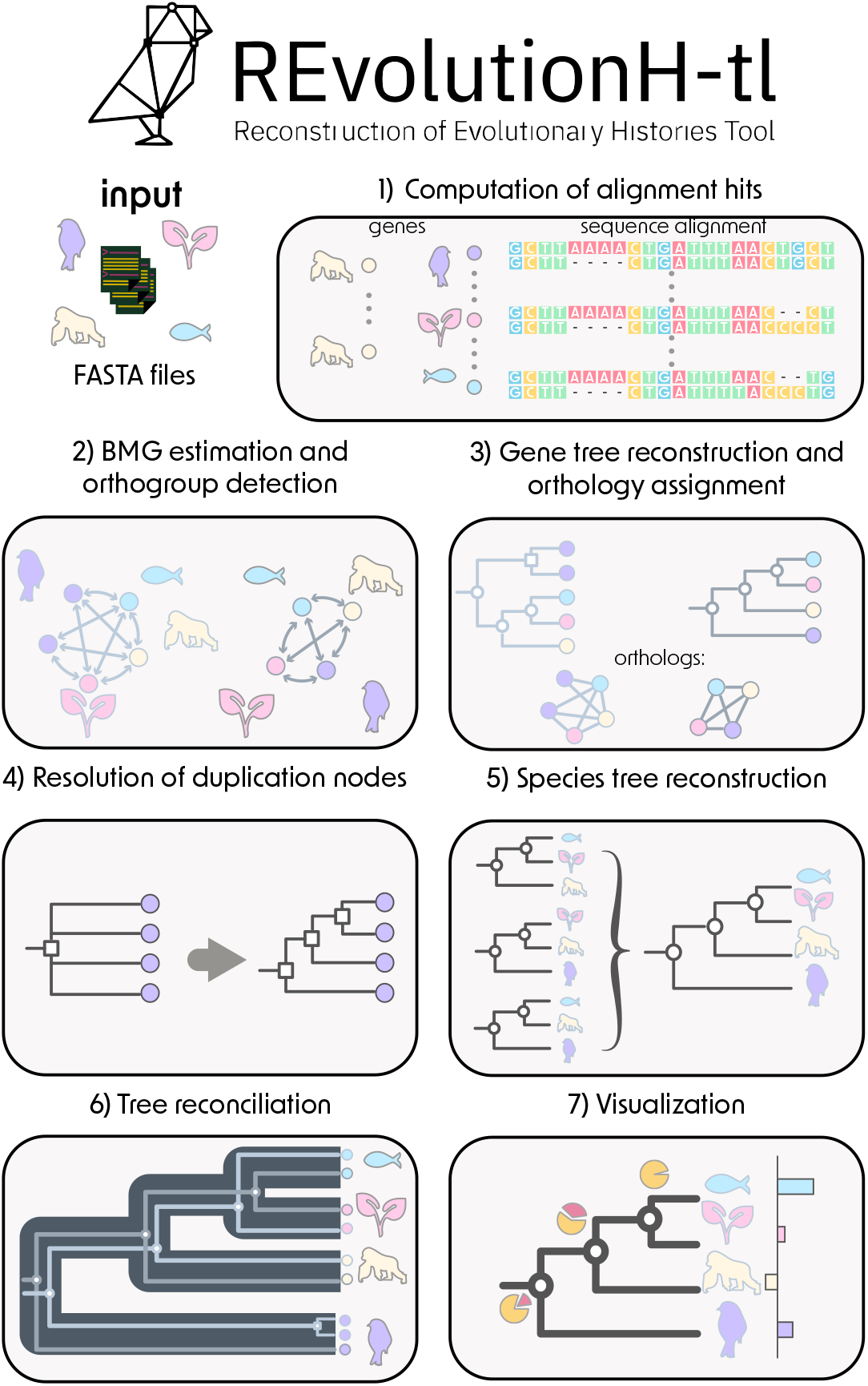
Conceptual overview of REvolutionH-tl. (1) DNA and protein alignments are computed using `BLAST` and `Diamond`. (2) BMGs are built from best hits; each connected component defines an orthogroup. (3) Gene trees are inferred per BMG, with duplication and speciation events assigned. Orthologs are identified when their last common ancestor is a speciation node. (4) Polytomic duplication nodes are resolved using a modified Neighbor-Joining algorithm. (5) A species tree is reconstructed from color triples derived from gene trees. (6) Gene trees are reconciled with the species tree to infer losses, duplications, and to refine gene tree structure. (7) Visualizations summarize reconciliations and report statistics on orthologs and orthogroups.

### Computation of Alignment Hits

The aim of this step is to compute sequence similarity between genes of different species. The input is a collection of fasta files, each of them corresponding to one species and containing a list of sequences. Alignments are computed using `Diamond` [42] for amino acid sequences or `BLAST` [43, 44] for DNA sequences. As output, `REvolutionH-tl` generates a directory containing the alignment hits for each pair of species.

### BMG Estimation and Orthogroup Detection

For a given gene *x* in species *A*, a gene *y* in species *B* is defined as a *best match* if it is evolutionarily the most closely related gene to *x* among all genes in species *B* [45]. Importantly, best matches are not necessarily unique.

If species *B* contains an ortholog *z* of *x*, then *z* is guaranteed to be a best match of *x*, assuming no horizontal gene transfer. This is because no gene in species *A* can be more closely related to a gene in *B* than gene pairs that arose from the speciation event separating *A* and *B*. However, the converse does not hold: even if an ortholog of *x* has been lost in *B, x* will still have a best match in *B*, provided that *B* contains any homolog of *x*.

The (typically asymmetric) directed graph formed by all such best match relationships is known as the *best match graph* (BMG).

In practice, best matches are often approximated by *best hits*, which are identified based on sequence similarity scores [16]. `REvolutionH-tl`follows this approach. However, it is important to distinguish between best hits and true best matches, as illustrated in Fig. 3.

**Fig 3.**
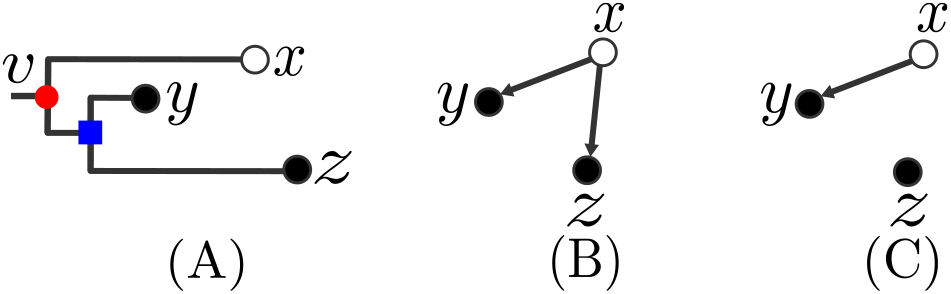
Best matches and best hits. (A) A gene tree with three genes *x, y*, and *z* from two species (represented in black and white), rooted at node *v*. The least common ancestor of *x* relative to *y* and *z* is *v*. The similarity score (measured as weighted tree distance) between *x* and *y* is higher than that between *x* and *z*, indicating that the Molecular Clock Hypothesis does not hold in this case. (B) Genes *y* and *z*, both from the black species, are best matches of *x*. (C) Gene *y* is the best hit of *x*, whereas *z*, though a best match, is not considered a best hit.

Specifically, `REvolutionH-tl`computes alignment hits between genes from different species and calculates a normalized bit score to approximate evolutionary relatedness. Following the method in [23], a gene *y* in species *B* is considered a best hit of *x* if its bit score falls within an adaptive threshold relative to the top-scoring hit.

This process produces a directed *best hit graph*, which reflects the sequence similarity relationships. The connected components of this graph are interpreted as *orthogroups*, representing sets of genes likely descended from a common ancestor.

The output of this step includes two `tsv` files: one listing the best hit gene pairs, and the other containing the identified orthogroups.

### Gene Tree Reconstruction and Orthology Assignment

Gene trees can be reconstructed from best match graphs (BMGs) by computing their so-called *least resolved trees* (LRTs) [45]. This process begins by extracting a set of *informative triples* from the best hit graph, following the approach described in [46]. However, because the best hit graph is only an approximation of the true BMG, the resulting triple set `ℛ` is typically inconsistent. To address this, `REvolutionH-tl`applies a heuristic to identify a maximum subset of consistent triples, denoted 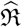 [46, 47].

The gene tree *T* is then reconstructed from 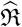 using the well-established `BUILD` algorithm [33]. Although the resulting tree is generally not fully resolved, it captures all orthology relationships among the genes in the corresponding orthogroup.

Next, internal nodes of the gene tree are annotated with evolutionary events. Each node *v* is labeled with an event type *t*(*v*) using the *species overlap* method [39, 48]. According to this method, *v* is labeled as a speciation event if the sets of species under its child subtrees do not overlap, and as a duplication event otherwise. The result is an event-labeled gene tree (*T, t, σ*), where *σ* maps each leaf (gene) to its corresponding species.

From the event-labeled tree, orthologous gene pairs are identified as those whose least common ancestor in *T* is labeled as a speciation event. A key strength of this relation-based approach is its robustness: correct orthology assignments require only the correct *discriminating tree*, which is obtained by collapsing consecutive internal nodes with the same event label. Thus, even a partially resolved gene tree with a few incorrect branches can yield accurate orthology predictions, as long as the overall structure captures the correct evolutionary splits.

This step produces three output files in `tsv` format: (1) a list of inferred gene trees in NHX format, each associated with its orthogroup ID; (2) a list of best matches recomputed from (*T, t, σ*); and (3) a list of orthologous gene pairs derived from the event-labeled tree.

### Polytomy Resolution for Gene Trees

Gene trees produced in the previous step are generally not fully binary. To resolve polytomies labeled as speciation events, we leverage the reconciliation map and the topology of the species tree. For polytomies labeled as duplication events, we apply a distance-based clustering strategy that incrementally builds a binary subtree using a neighbor joining (NJ) criterion, as described in Algorithm 1.

Let *x* `∈` *V* (*T*) be a duplication node in a gene tree *T*, with children ch(*x*) = {*y*_1_, *y*_2_, …, *y*_*k*_}, where *k* `≥` 3. The goal is to replace the unresolved polytomy at *x* with a fully resolved binary subtree *T* `′`.

We begin by initializing an unrooted star-shaped tree *T* `′`, whose leaf set is *L*(*T* `′`) = {*y*_1_, *y*_2_, …, *y*_*k*_}, and a central internal node *x*`′`. We also define a set of clusters *C* = {*C*_1_, *C*_2_, …, *C*_*k*_}, where each cluster *C*_*i*_ corresponds to the leaves of the subtree *T* (*y*_*i*_). For each pair *y*_*i*_, *y*_*j*_ `∈` ch(*x*), we compute a pairwise distance *d*(*y*_*i*_, *y*_*j*_) as the average of all distances *δ*(*z*_*i*_, *z*_*j*_) between gene pairs (*z*_*i*_, *z*_*j*_) `∈` *C*_*i*_ × *C*_*j*_. The distances *δ* are derived from sequence similarity scores obtained during the initial comparison step using `DIAMOND` or `BLAST`.

With the star tree *T* `′` and the distance matrix *d* in place, the algorithm proceeds iteratively. At each step, the pair *y*_*i*_, *y*_*j*_ `∈` *N* (*x*`′`) minimizing the NJ criterion is selected. These nodes are merged into a new internal node *y*_*ij*_: the edges *x*`′`*y*_*i*_ and *x*`′`*y*_*j*_ are removed, and new edges *x*`′`*y*_*ij*_, *y*_*ij*_*y*_*i*_, and *y*_*ij*_*y*_*j*_ are added. The distance matrix is updated using standard NJ rules to reflect distances between *y*_*ij*_ and the remaining neighbors of *x*`′`.

This iterative process continues until *T* `′` is fully resolved. A root is then added to *T* `′` using the midpoint rooting method [49]. We use the `Biopython` implementation of NJ and midpoint rooting algorithms [50].

The resulting binary subtree *T* `′` is then grafted back into the original gene tree *T*, replacing the unresolved duplication node *x*. Because the method relies solely on distance information and does not impose external biological constraints, it is particularly suitable for resolving duplication nodes in cases where topological refinement is needed independently of additional annotations.

#### Algorithm 1

Resolve-Duplication-Polytomy

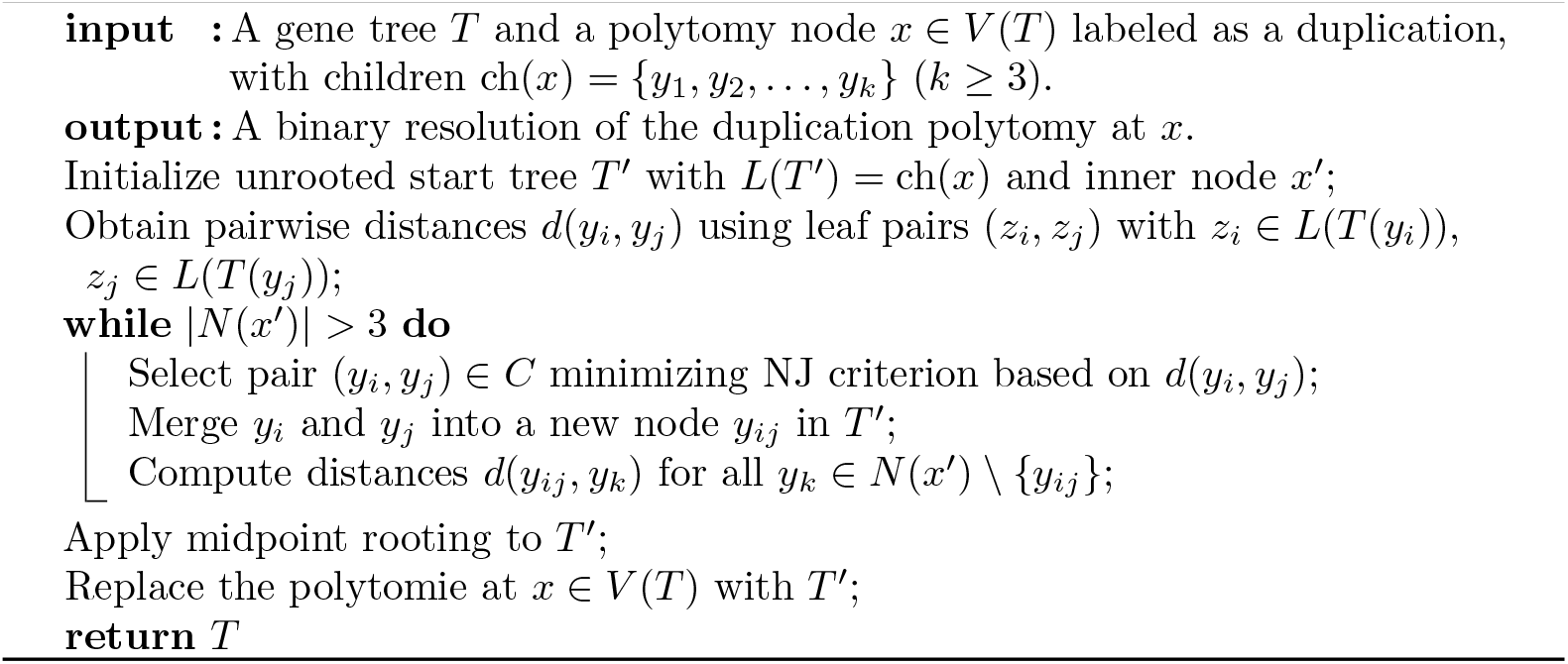

### Species Tree Reconstruction

Event-labeled gene trees (*T, t, σ*) contain implicit information about the underlying species tree *S* in the form of gene triples of the type ((*u, v*), *w*), where *u, v*, and *w* are genes from three different species. If their lowest common ancestor lca(*u, w*) = lca(*v, w*) is labeled as a speciation event, then the corresponding species triple ((*σ*(*u*), *σ*(*v*)), *σ*(*w*)) is displayed by the species tree [31, 32, 51].

In the absence of noise, reconstructing the species tree from these triples can be accomplished by applying the `BUILD` algorithm to the set of inferred species triples. However, to account for errors in the input data—particularly those introduced in earlier processing steps—a heuristic extension of `BUILD` is employed, as proposed in [46]. This heuristic modifies the behavior of `BUILD` by forcing a partition of the auxiliary Aho graph whenever it remains connected, ensuring progress in the reconstruction even in the presence of conflicts or inconsistencies.

The output of this step is a single file containing the inferred species tree in NHX format.

### Tree Reconciliation

An *evolutionary scenario* is defined as a mapping *µ* from a gene tree (*T, t, σ*) into a species tree *S*, satisfying the following conditions: (i) each gene tree leaf *x* `∈` *L*(*T*) is mapped to the corresponding species leaf *µ*(*x*) = *σ*(*x*); (ii) each internal node of *T* labeled as a speciation is mapped to an internal node of *S*; (iii) each duplication node in *T* is mapped to an edge in *S*; and (iv) the mapping *µ* preserves the ancestor-descendant relationships, i.e., if *u* is an ancestor of *v* in *T*, then *µ*(*u*) is an ancestor of *µ*(*v*) in *S* (see Fig. 1).

Before computing the reconciliation map, we first assess whether the gene tree and the species tree are topologically consistent. This is achieved by verifying that all species triples implied by the gene tree’s color-labeled triples are also present in the species tree. If this condition is not met, we apply a correction procedure to the gene tree. Specifically, we identify leaves involved in the most inconsistent and least consistent triples and prune them, as detailed in Algorithm 2.

Once the gene tree has been corrected and is consistent with the species tree, reconciliation is performed by mapping each node of the gene tree to a node or edge in the species tree according to its evolutionary label. Leaf nodes are mapped directly to their corresponding species. Internal nodes labeled as speciations are mapped to internal nodes in the species tree, while duplication nodes are mapped to edges, preserving the topological order of divergence events.

#### Algorithm 2

Prune-L: Correction of inconsistent gene trees

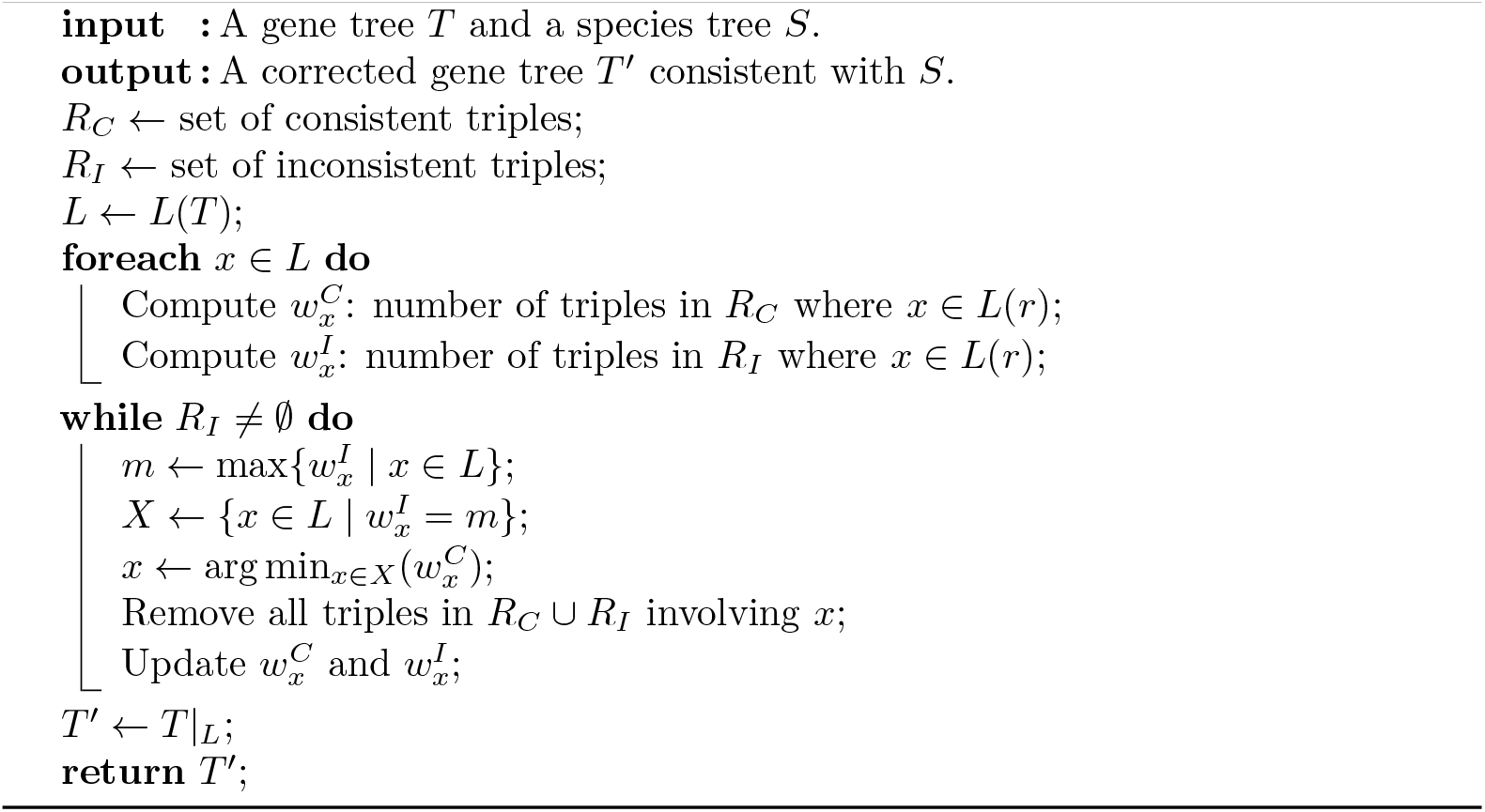

The corrected gene tree may still contain unresolved polytomies. For nodes labeled as speciations, these will be resolved during reconciliation, guided by the topology of the species tree. Duplication nodes are resolved applying the modified NJ algorithm described above.

Gene loss events are inferred during reconciliation by examining speciation nodes in the gene tree. When a speciation node *u* is mapped to an internal node *v* in the species tree, we verify whether each child of *v* is represented among the descendants of *u*. If this is not the case, gene loss events are inferred accordingly.

The input for this step consists of a list of gene trees and a species tree, both in NHX format. The user may provide a species tree, or it can be inferred automatically by `REvolutionH-tl`as described in the previous section. The output is a `tsv` file containing evolutionary scenarios. Each entry includes the orthogroup identifier, the reconciled gene tree in NHX format, and a reconciliation map that associates each node in the gene tree with its corresponding node or edge in the species tree. Since only duplication nodes are mapped to edges, we report the child node *v* of the edge *uv* in *S* as the reconciliation target for a duplication node. As a result, the reconciliation map is presented as a list of node pairs. The reconciled gene trees produced by this step explicitly annotate evolutionary events, including gene duplications and losses.

### Visualization of Reconciliation Results

We provide dedicated commands for visualizing the reconciliation results. The command revolutionhtl.plot reconciliation allows users to display the evolutionary scenario of a specific orthogroup by embedding the gene tree within the species tree. This visualization facilitates intuitive interpretation of the gene-species mapping and the associated evolutionary events. An example is shown in Fig. 7.

For a broader summary across multiple gene families, the command revolutionhtl.plot summary generates two comprehensive diagrams: (i) the *reconciliation tree*, and (ii) a diagram illustrating the *relative change in gene content* across clades.

The reconciliation tree provides an overview of gene content evolution across the species tree. It reports counts of inherited genes, duplicated genes, clade-specific gains, and species-specific genes, as illustrated in Fig. 4. Additionally, it summarizes orthogroup statistics, including the number of single-copy orthogroups, the average number of species per orthogroup, classifications of orthology relationships, and the total number of single-copy genes. A representative example is shown in Fig. 5.

**Fig 4.**
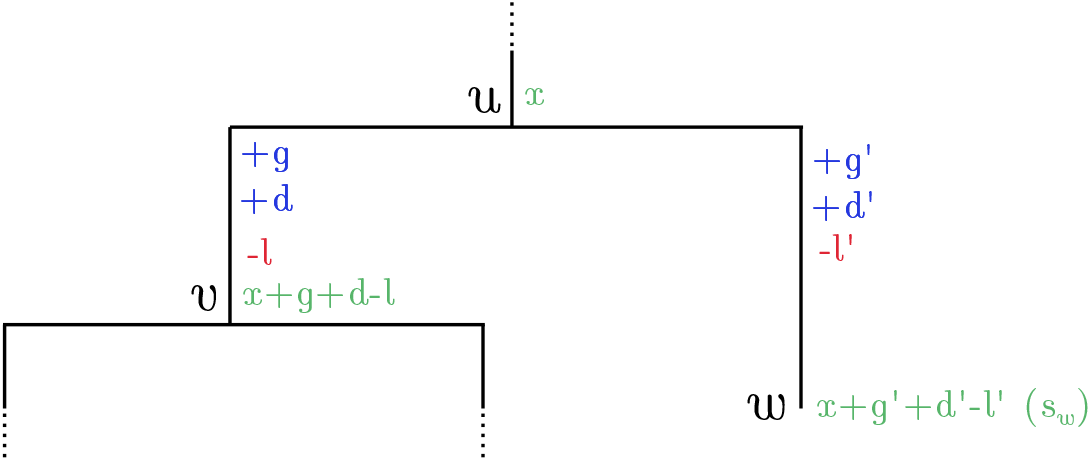
Explaining change in gene content. We show two inner nodes and one leaf of a species tree, labeled as *u, v*, and *w* respectively. Additionally, we add some integer variables explaining the evolution of gene content for these three elements. For the inner nodes *u* and *v*, the green numbers correspond to genes present in a species just before the speciation event, for example, node *u* has *x* genes while *v* has *x* + *g* + *d* `−` *l*. Numbers along the branch *uv* display gene acquisitions and losses in color blue and red correspondingly; *g* is the number of gained genes (corresponding to the number of gene families specific for *v* and descendants) while *d* stands for the number of duplicated genes, and the red number *l* corresponds to gene loss. The gene content of leaf *w* is explained similarly, we just add an extra variable in parenthesis, *s*_*w*_, which shows the number of species-specific genes, also known as singletons.

**Fig 5.**
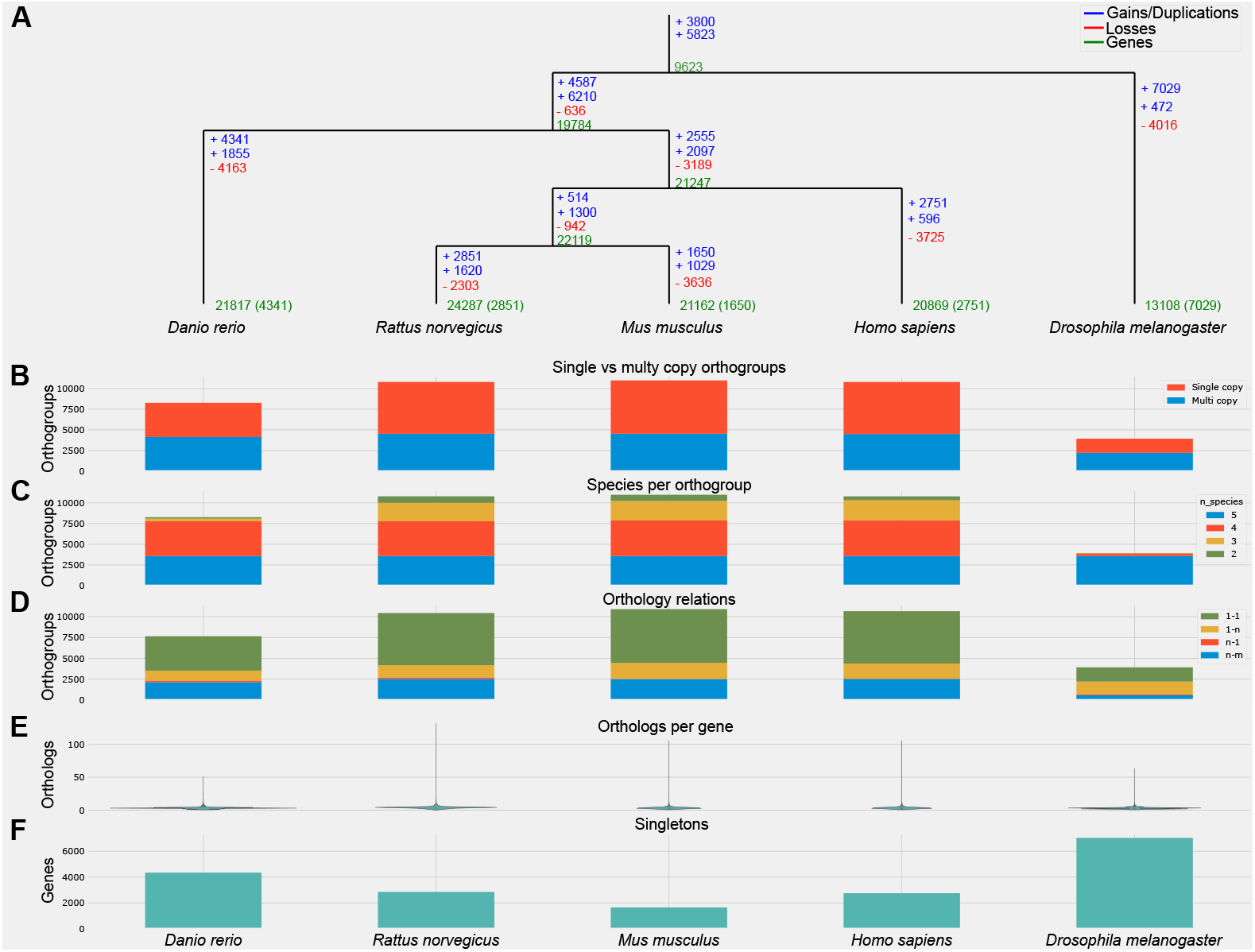
Genome complexity evolution for the Phylome05 dataset. The species tree shown at the top of the figure defines the left-to-right order of species along the x-axis: *Danio rerio, Rattus norvegicus, Mus musculus, Homo sapiens*, and *Drosophila melanogaster*. Each column in panels (B–F) corresponds to one of these species, as ordered by the tree, and displays species-specific data. For example, the column below *Danio rerio* exclusively reports statistics derived from orthogroups that contain at least one gene from this species. (A) Evolution of genome complexity, following the notation introduced in Fig. 4. Gene counts per species are shown in green, while blue and red indicate gene gains/duplications and gene losses, respectively. (B) Proportion of orthogroups consisting of single-copy genes versus those containing paralogs. (C) Distribution of orthogroups based on the number of species represented within each group. (D) Classification of orthology relationships for genes from the species associated with the current column. Relationships are categorized as one-to-one, one-to-many, many-to-one, or many-to-many across species. (E) Number of orthologs per gene for the species indicated by the column. (F) Total count of species-specific genes. Font sizes in the figure were manually adjusted to enhance readability.

The second diagram highlights clades undergoing the most significant shifts in gene content. It reports the percentage of ancestral versus *de novo* duplications and applies min-max normalization to the metrics reported in Fig. 4. This visualization enables quick identification of clades with particularly dynamic evolutionary histories. An example is shown in Fig. 6.

**Fig 6.**
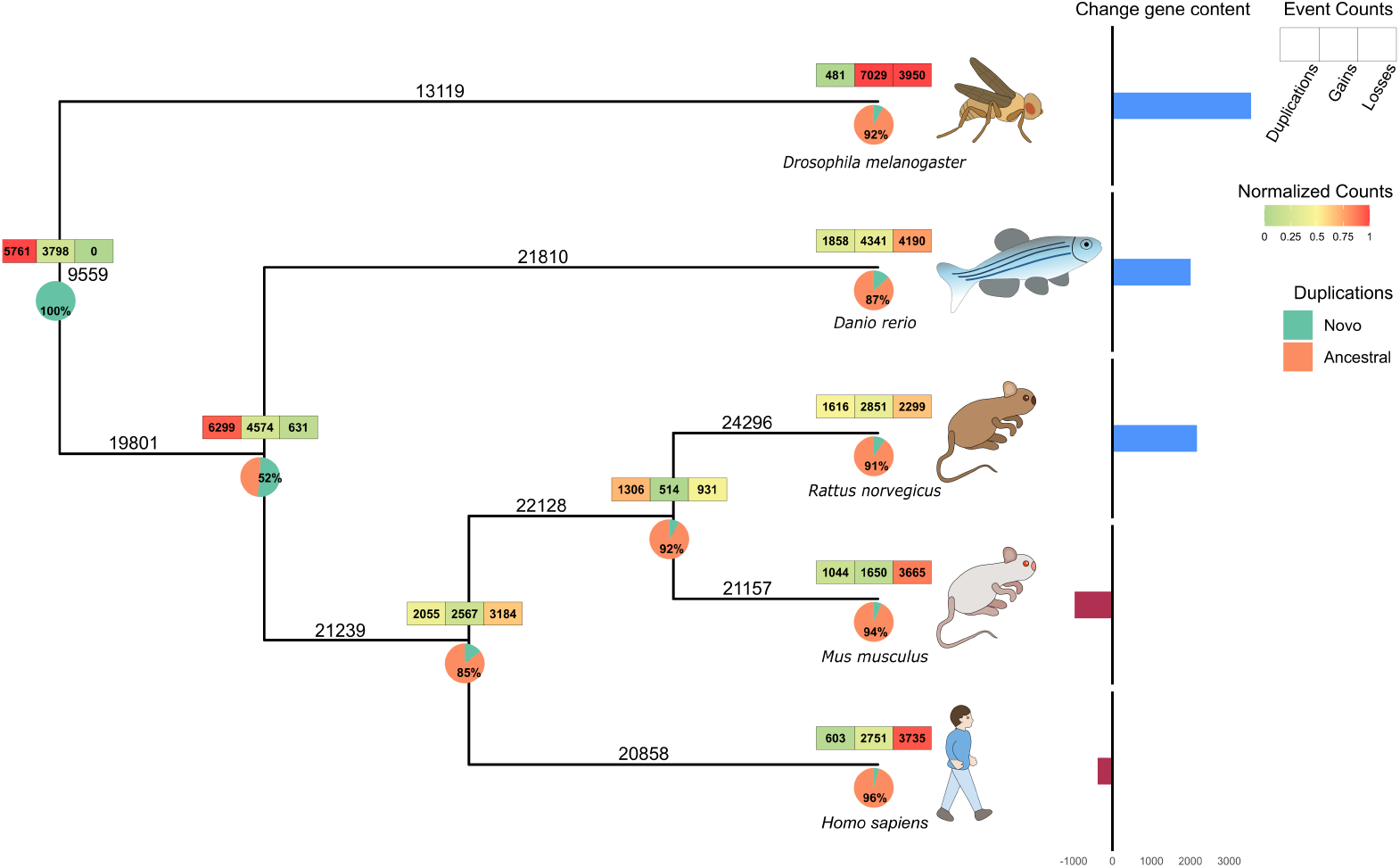
Visualization of relative change in gene content for the Phylome0500 dataset using REvolutionH-tl. Summary of key evolutionary events across the phylogeny, including gene duplications, gains, and losses. A heatmap along the branches encodes normalized event values, where red indicates below-average and green indicates above-average frequencies. At each internal node, pie charts represent the proportion of *de novo* versus ancestral gene duplications. To the right of the tree, bar charts depict changes in gene content across species: blue bars indicate gene gains, while wine-colored bars correspond to gene losses. Visual representations of the analyzed species were added manually to enhance interpretability.

**Fig 7.**
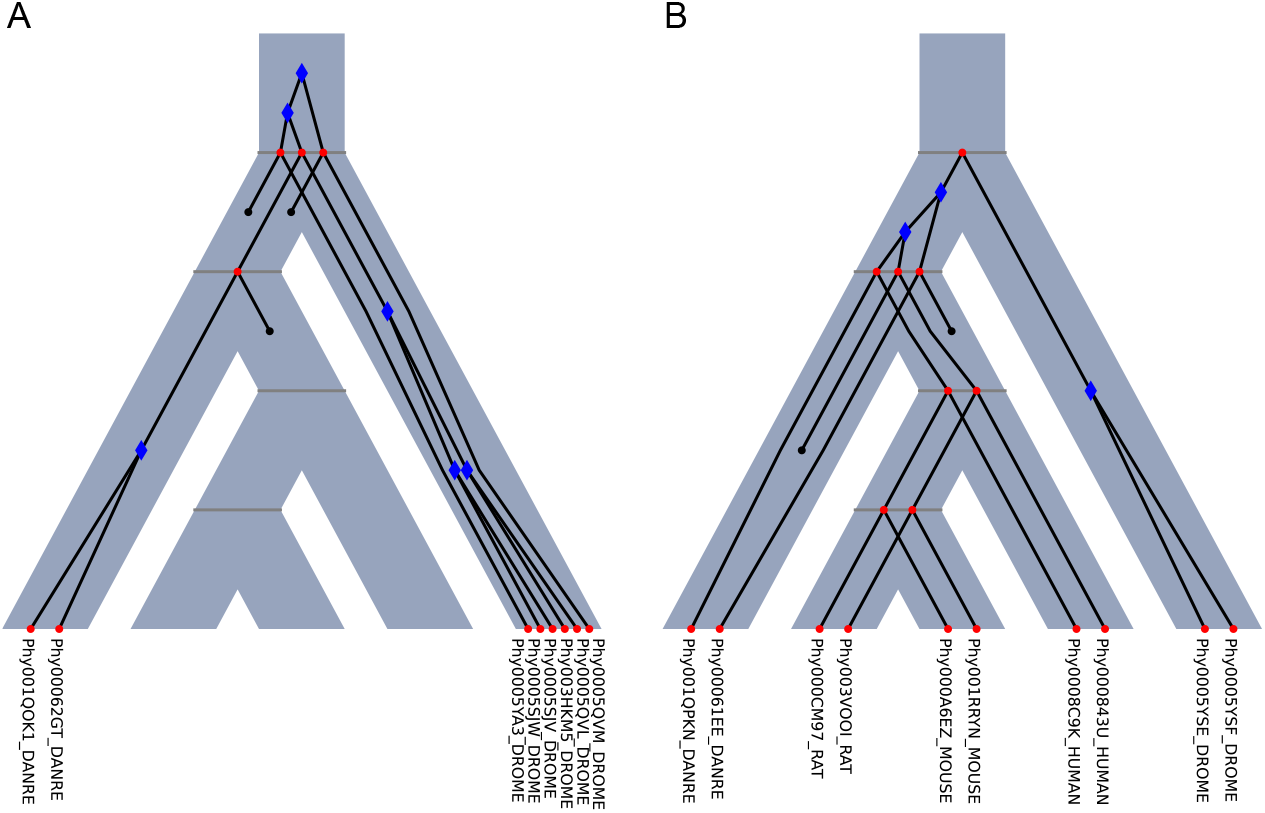
Evolutionary scenarios from the Phylome05 dataset. The species tree is represented as a gray pipe, within which the gene trees evolve. Gene trees are overlaid in black, with bifurcations annotated to indicate evolutionary events. Divergences caused by speciation are marked with a red 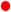, while gene duplication events are indicated with a blue 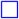. Additionally, gene tree leaves are color-coded: red denotes extant genes, whereas black signifies gene loss events.

All visualizations are generated using custom R scripts (v4.4.0) integrated into the `REvolutionH-tl`framework. The layout and design were inspired by visualization strategies proposed in [52].

## Benchmarking

To show the correctness of our approach, we run the `REvolutionH-tl`workflow to predict (i) orthogroups, (ii) orthology, (iii) gene trees, (iv) species trees, and (v) tree reconciliation. Then, we contrasted these predictions against those made by `Proteinortho`, `Orthofinder`, `RANGER-DTL`, `RAxML`, `GeneRax`, and `ASTRAL-Pro`. Table 1 illustrates how we compare the outputs of revolutionhtl and the other tools.

**Table 1.** Input and output of the benchmarking tools. Rows of this table corresponds to the different tools used for the benchmarking analysis. On the other hand, columns of the table show input/output files. The checkmark (✓) indicates the outputs that we keep for each took, similarly, the letter *I* indicates the input files.

|  | Molecular Model | MSA | genes/proteins | Orthogroups | Orthology | Gene Trees | Species Trees | Reconciliation |
| --- | --- | --- | --- | --- | --- | --- | --- | --- |
| REvolutionH-tl |  |  | I | ✓ | ✓ | ✓ | ✓ | ✓ |
| Proteinortho |  |  | I | ✓ | ✓ |  |  |  |
| Orthofinder |  |  | I | ✓ | ✓ |  |  |  |
| RAxML | I | I |  |  |  | ✓ |  |  |
| GeneRax | I | I |  |  |  |  | I | ✓ |
| Astral-Pro |  |  |  |  |  | I | ✓ |  |
| RANGER-DTL |  |  |  |  |  | I | I | ✓ |

### Synthetic Dataset

We use `SaGePhy` [9] to simulate evolutionary histories under three main parameters: the number of species *N* `∈` {5, 10, 15, …, 50}, duplication rate *d*, and loss rate *l*. The parameters *d* and *l* are independently and uniformly sampled from the set {*i/*10 | *i* `∈ N`, 0 `≤` *i* `≤` 10}, resulting in a total of 1,210 distinct parameter combinations (*N, d, l*).

For each parameter triple, we simulate five independent evolutionary scenarios on a species tree *S*, resulting in five gene trees that evolve within *S*. The simulated trees include branch lengths, which reflect evolutionary rates under the assumption of a constant mutation rate across all genes. Protein sequences are then generated based on these trees using the *LG substitution model* [53], one of the models provided by `SaGePhy`.

To construct genome-like datasets, we extract the simulated protein sequences and generate a set *F*_*N*_ of FASTA files, one per species. Each FASTA file represents the complete proteome of a simulated species. We refer to the resulting dataset of simulated genomes and evolutionary histories as `SaGePhy5-50`.

This synthetic dataset provides a controlled ground truth for evaluating inference tools. Because the evolutionary scenarios are fully known, we can directly extract the *true* orthogroups and orthology graphs. Performance evaluation is then conducted by comparing these ground truth annotations to the predictions produced by each tool.

### Metazoan Genomes

To demonstrate the application of our tool on real biological data, we selected five species from the `(PhyId 500) [QfO] Reference Model Species Metaphylome` dataset available in the PhylomeDB database [48]. The selected species are: *Homo sapiens, Mus musculus, Rattus norvegicus, Danio rerio*, and *Drosophila melanogaster*.

For each species, we obtained FASTA files containing protein sequences. To ensure a consistent representation of gene loci, we excluded protein variants and isoforms, retaining only a single representative sequence per gene. It is important to note that, because we use proteome data, the sequences in these FASTA files correspond exclusively to protein-coding genes.

We refer to this dataset as `Phylome05`, and use it to evaluate the behavior and outputs of our pipeline on high-quality, curated genomic data.

### Correctness of Orthogroup Identification

Accurately identifying homologous genes is a fundamental step in evolutionary analysis. To evaluate the correctness of orthogroup predictions on the synthetic dataset, we compare the results produced by `REvolutionH-tl`, `ProteinOrtho`, and `Orthofinder` against the ground-truth orthogroups generated using `SaGePhy`.

To assess performance, we classify each ground-truth orthogroup according to the relationship between its true members and the predicted orthogroups, as illustrated in Fig. 8. A ground-truth orthogroup *X* is considered *fully recovered* if there exists a predicted orthogroup *X*`′` such that *X* = *X*`′`. If no such match exists, the prediction is deemed incorrect.

**Fig 8.**
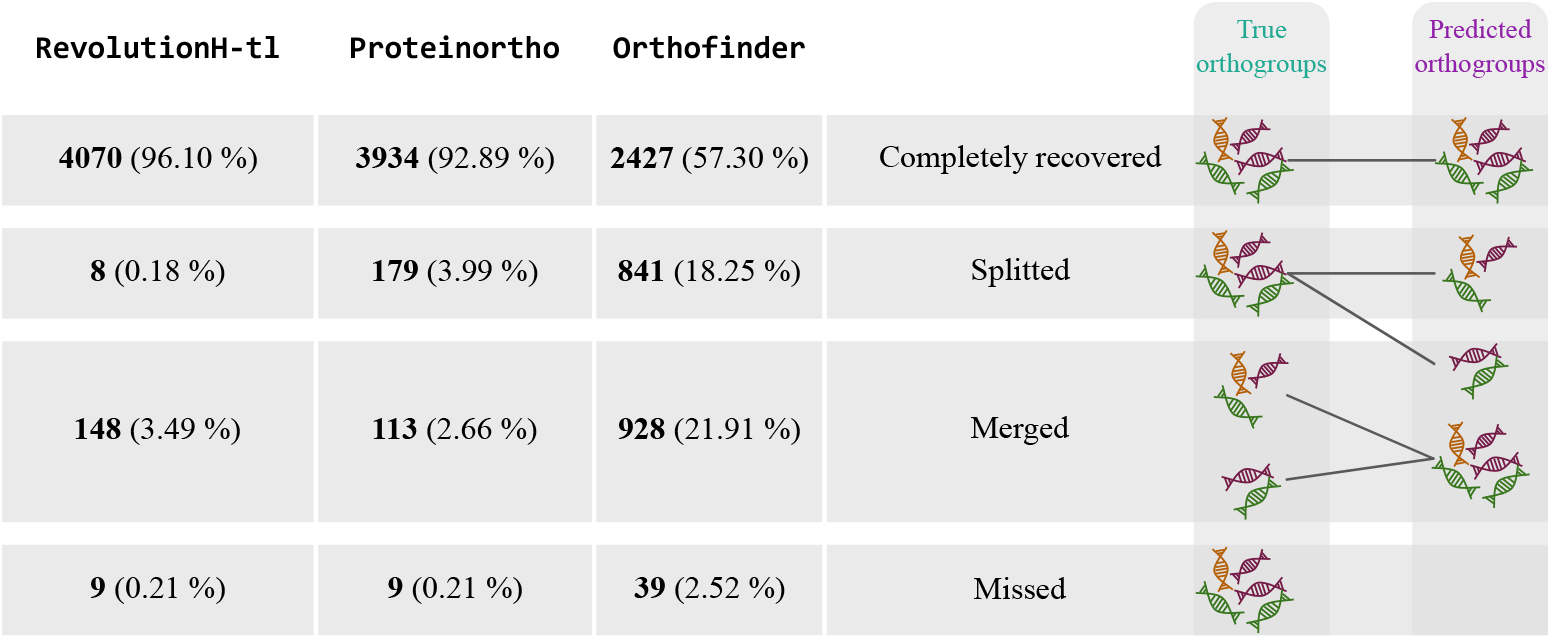
Orthogroup performance. Fully recovered orthogroups are those that exactly match a corresponding inferred orthogroup. Split orthogroups are those whose genes are distributed across multiple inferred orthogroups, indicating under-clustering. In contrast, merged orthogroups are entirely contained within a single inferred orthogroup that also includes genes from other true orthogroups, reflecting over-clustering.

Incorrect predictions are further categorized into two error types:

- Split orthogroups: A true orthogroup *X* is considered split if the predicted orthogroup is a proper subset, i.e., *X*`′ ⊂` *X*. This indicates that the tool failed to recover all members of the group in a single cluster. sets.
- Merged orthogroups: Two or more true orthogroups (e.g., *X*_0_ and *X*_1_) are considered merged if a predicted orthogroup *X*`′` contains elements from both, i.e., *X*`′ ∩` *X*_0_ ≠ ∅ and *X*`′ ∩` *X*_1_ ≠ ∅. This suggests an over-clustering of unrelated gene

For each tool, we report the percentage of correctly recovered, split, and merged orthogroups to quantify its performance in orthogroup identification.

### Standard Performance Metrics

To evaluate the performance of tools for orthology prediction and gene tree inference, each individual prediction is classified as a *true positive* (TP), *false positive* (FP), *true negative* (TN), or *false negative* (FN). These predictions may correspond to orthology relationships or resolved triplets in a gene or species tree. Once the predictions have been categorized, we compute the standard performance metrics as follows. Precision is defined as 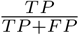, measuring the proportion of correct positive predictions. Recall is given by 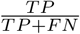, representing the proportion of actual positives that are correctly identified. The false positive rate (FPR) is computed as 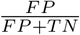, indicating the proportion of negatives incorrectly classified as positives. Finally, accuracy is calculated as 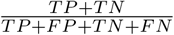, reflecting the overall proportion of correct predictions.

For each orthogroup *X*, we calculate its performance value *A*_*X*_ according to each of the metrics described. To assess trends under specific evolutionary conditions, orthogroups are grouped into collections based on shared simulation parameters, such as fixed duplication and loss rates. For a collection of orthogroups *Y* = {*X*_0_, *X*_1_, …}, the average performance for a given metric is computed as:

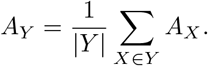

This aggregated evaluation enables a consistent and robust comparison of inference tools across diverse evolutionary scenarios.

### Performance for Orthology Inference

We evaluate the accuracy of orthology prediction at the level of individual orthogroups. Let *X* be a true orthogroup, defined as a set of genes. The set of true orthology relations within *X* is denoted by Θ_*X*_ `⊆` {*uv* `∈` *X* × *X* | *u* ≠ *v*}.

Let *I*_*X*_ {*uv* `∈` *X* × *X*`′` | *u* ≠ *v*} be the set of orthology relations inferred by a method, where *X* `⊆`*X*`′` and *X*`′` may include genes that do not belong to the true orthogroup *X*. This formulation allows the detection of both over-clustering and under-clustering errors.

We classify the outcomes of the predictions by computing the number of true positives as |*I*_*X*_ \Θ_*X*_ |, the number of false positives as |*I*_*X*_ `∩` Θ_*X*_ |, and the number of false negatives as |Θ_*X*_ \ *I*_*X*_ |. The number of true negatives is given by 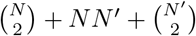, where *N* = |*X*| and *N* `′` = |*X*`′` \ *X*|, thus accounting for all pairwise gene combinations, including those within the orthogroup, between orthogroup and external genes, and among external genes themselves.

Based on these quantities, we compute standard performance metrics such as precision, recall, accuracy, and false positive rate.

### Performance for Tree Reconstruction

To assess the accuracy of inferred phylogenies, we compare each reconstructed tree *T* `′` with its corresponding ground-truth tree *T* by evaluating the sets of rooted triples they induce. A rooted triple represents the resolved relationship among a triplet of taxa in the tree.

Let *R*(*T*) and *R*(*T* `′`) be the sets of rooted triples derived from the true and inferred trees, respectively. From these, we calculate the number of true positives as |*R*(*T*) `∩` *R*(*T* `′`)|, the number of false positives as |*R*(*T* `′`) \ *R*(*T*)|, and the number of false negatives as |*R*(*T*) \ *R*(*T* `′`)|. The number of true negatives is computed as 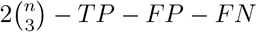, where *n* denotes the number of taxa.

These values form the basis for standard performance metrics, allowing us to evaluate the quality of tree reconstruction in terms of precision, recall, and false positive rate, relative to the resolved triplets in the true topology.

### Reconciliation Distance

To assess the quality of gene tree reconstructions, we reconciled the event-labeled gene trees inferred by all methods with the true species tree. This enabled us to apply the *PLR* dissimilarity measure [54], which estimates the confidence in reconciliation results by comparing evolutionary scenarios. Given two evolutionary scenarios, 𝒢_1_ = (*S, T*_1_, *t*_1_, *σ*_1_, *µ*_1_) and 𝒢_2_ = (*S, T*_2_, *t*_2_, *σ*_2_, *µ*_2_), the *PLR* dissimilarity is defined as:

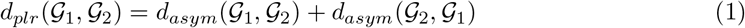

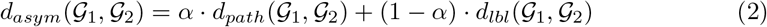

Equation 1 ensures symmetry, i.e., *d*_*plr*_(𝒢_1_, 𝒢_2_) = *d*_*plr*_(𝒢_2_, 𝒢_1_), while Equation 2 shows that the dissimilarity is composed of two components: the *path* component and the *label* (*lbl*) component.

Both components rely on a node-mapping function *m*, which maps each node *x* `∈` *V* (*T*_1_) to a corresponding node *y* `∈` *V* (*T*_2_), defined as 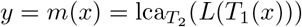. The *path* component measures the number of edges between *µ*_1_(*x*) and *µ*_2_(*y*) in the species tree *S*, capturing the topological distance between the mapped positions. The *lbl* component reflects differences in the evolutionary event labels of *x* and *y*, increasing by one if *t*_1_(*x*) ≠ *t*_2_(*y*), and zero otherwise.

In our experiments, we set *α* = 1*/n*, as recommended in [54], where *n* is the number of species in the species tree.

This reconciliation-based measure allows us to simultaneously evaluate structural differences between gene trees and inconsistencies in the predicted reconciliation mappings and evolutionary events, providing a comprehensive and interpretable metric for assessing gene tree accuracy.

## Results

Our primary contribution is `REvolutionH-tl`, an open-source, cross-platform Python tool [55] developed for fast and accurate orthology prediction, as well as the inference and reconciliation of gene and species trees. The design and functionality of `REvolutionH-tl`make it particularly well-suited for large-scale comparative genomics analyses.

Since its initial release [22], we have introduced several enhancements to both the software and the accompanying benchmarking framework to further validate the correctness of our inferences. Methodological improvements include the integration of a novel module for polytomy resolution. This module complements the existing best-match strategy by enabling the binarization of polytomies labeled as duplication events, thereby improving the biological realism of the inferred gene trees.

In addition to methodological advancements, we have expanded the software’s visualization capabilities. New commands now allow for the graphical exploration of evolutionary scenarios, facilitating integrative and interpretable analyses. Furthermore, runtime performance has been significantly improved through the parallelization of key modules responsible for processing alignment hits and selecting best matches.

To evaluate the performance of `REvolutionH-tl`, we conducted an extended benchmarking study that includes six widely used tools for evolutionary analysis: `Proteinortho`, `Orthofinder`, `RAxML`, `Astral-Pro`, `RANGER-DTL`, and `GeneRax`. For each of these tools, we used the latest available version to ensure a fair and up-to-date comparison.

### Evolution Visualization

We provide the command revolutionhtl.plot summary for the comprehensive aggregation and visualization of the evolutionary history of multiple gene families. This functionality is exemplified in Figures 5 and 6, which illustrate the evolution of genome complexity across metazoan species included in the `Phylome05` dataset. The evolutionary scenarios for this analysis were inferred directly from FASTA files using the command python -m revolutionhtl -F fastas/.

In addition, this plotting command accepts parameters that allow users to focus on a custom subset of orthogroups, enabling targeted analyses of specific gene families of interest.

Additionally, the command revolutionhtl.plot reconciliation enables the visual inspection of complex evolutionary scenarios by embedding gene trees within the species tree context. In these visualizations, the species tree is represented as a gray pipe along which genes evolve. Gene trees are drawn in black, with bifurcations annotated to indicate evolutionary events: red 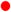 marks speciation events, and blue 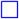 denotes gene duplications. Gene tree leaves are also color-coded—red for extant genes and black for inferred gene losses.

The command revolutionhtl.plot reconciliation enables the visual inspection of complex evolutionary scenarios by embedding gene trees within the species tree context. In these visualizations, the species tree is represented as a gray pipe along which genes evolve. Gene trees are drawn in black, with bifurcations annotated to indicate evolutionary events: red 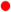 marks speciation events, and blue 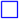 denotes gene duplications. Gene tree leaves are also color-coded—red for extant genes and black for inferred gene losses.

For example, Fig. 7A illustrates a gene family with multiple duplication events, both ancestral and species-specific, as well as three gene losses. This results in a family composed of only eight genes distributed across two species. In contrast, Fig. 7B shows a more conserved scenario in which the gene family is retained across all sampled species. These visualizations facilitate the interpretation of evolutionary dynamics by providing an intuitive depiction of gene gain, duplication, and loss events within the broader species tree framework.

#### Precise Orthogroups

Figure 8 compares the performance of `REvolutionH-tl`, `Orthofinder`, and `Proteinortho`in identifying orthogroups from the dataset `SaGePhy5-50`. Our results show that `REvolutionH-tl`achieves the highest accuracy, with nearly all inferred orthogroups precisely matching their corresponding ground-truth orthogroups. In comparison, `Proteinortho`exhibits a moderate reduction in accuracy, recovering approximately 3% fewer orthogroups, while `Orthofinder`shows a substantial performance decline, recovering nearly 40% fewer orthogroups than the ground truth.

To further investigate the sources of error, we analyze orthogroup prediction failures by quantifying two error types: orthogroups that are split across multiple predicted clusters (i.e., split orthogroups), and those that are incorrectly grouped together into a single cluster (i.e., merged orthogroups). Across all tools, merging emerges as the predominant source of error. Notably, `Orthofinder`merges nearly 30% of the true orthogroups.

### High Accuracy of Orthology Prediction

Figure 9A presents the distribution of orthology prediction accuracy for `Orthofinder`, `Proteinortho`, and `REvolutionH-tl`on the `SaGePhy5-50` dataset. As expected, all tools exhibit high accuracy, owing to the ultrametric nature of the simulated dataset. Among them, `Proteinortho`achieves slightly higher accuracy than `REvolutionH-tl`in recovering orthologous gene pairs, while `Orthofinder`demonstrates a significantly lower accuracy in orthology inference.

**Fig 9.**
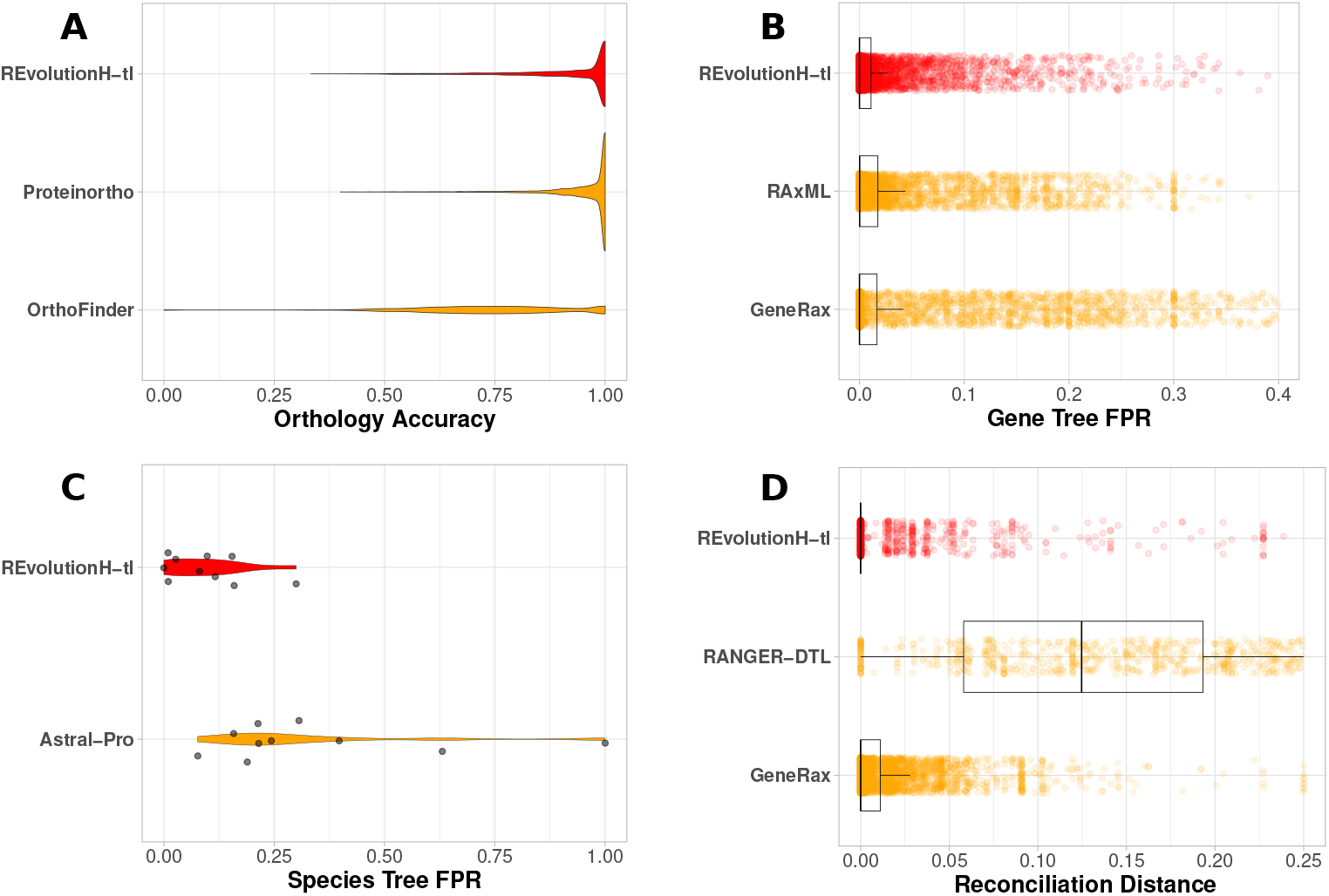
Benchmarking results for orthology prediction, gene and species tree reconstruction, and reconciliation. (A) Orthology accuracy. Each tool was evaluated on the same input: a collection of FASTA files from the `SaGePhy5-50` dataset. All runs were performed using 16 CPUs. For `Orthofinder`, the -M dendroblast parameter was used; default settings were applied for both `REvolutionH-tl`and `Proteinortho`. (B) Gene tree false positive rate (FPR). FPR distributions are computed from rooted triples obtained from inferred gene trees using the `SaGePhy5-50` dataset. `REvolutionH-tl`triples are derived from 5,736 fully recovered gene trees following reconciliation. `RaxML` triples are based on 5,848 inferred trees, and `GeneRax` triples on 5,885 trees. For `RAxML`and `GeneRax`, input consists of multiple sequence alignments (MSAs) generated with `MAFFT`. Both tools also require a molecular evolution model; we used the NJ-based model configuration for consistency. (C) Species tree false positive rate. This panel shows the FPR of rooted triples from species trees inferred from the `SaGePhy5-50` dataset. Each dot represents a species tree with `{`5, 10, …, 35, 40, 50} taxa, inferred from up to 600 gene trees. (D) Reconciliation distance. Distances are calculated using the normalized PLR dissimilarity measure between the inferred evolutionary scenarios and the ground-truth histories from the `SaGePhy5-50` dataset. `GeneRax`reconciled 5,885 gene trees, while `REvolutionH-tl` reconciled 5,736 gene trees, of which 3,341 are fully displayed without additional pruning. The same set of orthogroups was used as input for `RANGER-DTL`, which reconciles the gene trees produced by `REvolutionH-tl`at step 3.

This performance trend remains consistent when results are stratified by the simulation parameters used to generate the evolutionary scenarios, particularly the rates of gene duplication and loss. For a more comprehensive analysis of orthology prediction under varying evolutionary conditions, we refer the reader to [22].

### Accuracy for Gene and Species Tree Reconstruction

Figure 9B compares the distribution of false positive rates (FPR) for gene tree triples produced by three methods: `REvolutionH-tl`, `RaxML`, and `GeneRax`. All three tools maintain a low FPR, indicating reliable performance in capturing true topological signals. Notably, both `RaxML` and `GeneRax` rely on maximum likelihood methods and use the LG substitution model - the same model applied to simulate sequences in the `SaGePhy5-50` data set. Based on this methodological alignment, one might expect these tools to outperform `REvolutionH-tl`. However, it is noteworthy that `REvolutionH-tl` also achieves a comparably low FPR, despite relying on a simpler and faster approach.

Figure 9C shows the FPR distributions for species tree triples inferred by `REvolutionH-tl`and `Astral-Pro`on the `SaGePhy5-50` dataset. For each number of species in the set {5, 10, 15, …, 35, 40, 50}, approximately 600 gene trees are used to infer a single species tree. In this comparison, `REvolutionH-tl`operates on partially resolved gene trees, whereas `Astral-Pro`introduces random binary resolutions for all polytomies. This difference in treatment of unresolved nodes likely contributes to the superior accuracy of species trees inferred by `REvolutionH-tl`, as it avoids introducing potentially misleading topological artifacts through arbitrary resolutions.

### Precise Tree Reconciliation

Figure 9D presents the distribution of reconciliation distances for `REvolutionH-tl`, `RANGER-DTL`, and `GeneRax`, evaluated on the `SaGePhy5-50` dataset. All three tools demonstrate the ability to reconcile gene trees with the true species tree to a reasonable extent. However, the distribution for `REvolutionH-tl`is notably concentrated near zero, indicating that the evolutionary scenarios it produces are, on average, closer to the ground-truth evolutionary history. In contrast, the reconciliations generated by `GeneRax` and `RANGER-DTL` exhibit higher variability and larger distances, suggesting greater divergence from the simulated evolutionary scenarios.

### Fast Reconstruction of Evolution

The runtime performance of the tools included in our benchmarking analysis is shown in Fig. 10A. `REvolutionH-tl`successfully completes the full analysis of genomes from the `SaGePhy5-50` dataset. Notably, even for the largest dataset, `REvolutionH-tl`completes the analysis in under one hour. This represents a substantial improvement over the previous release of the tool, where the same analysis required several hours to complete [22].

**Fig 10.**
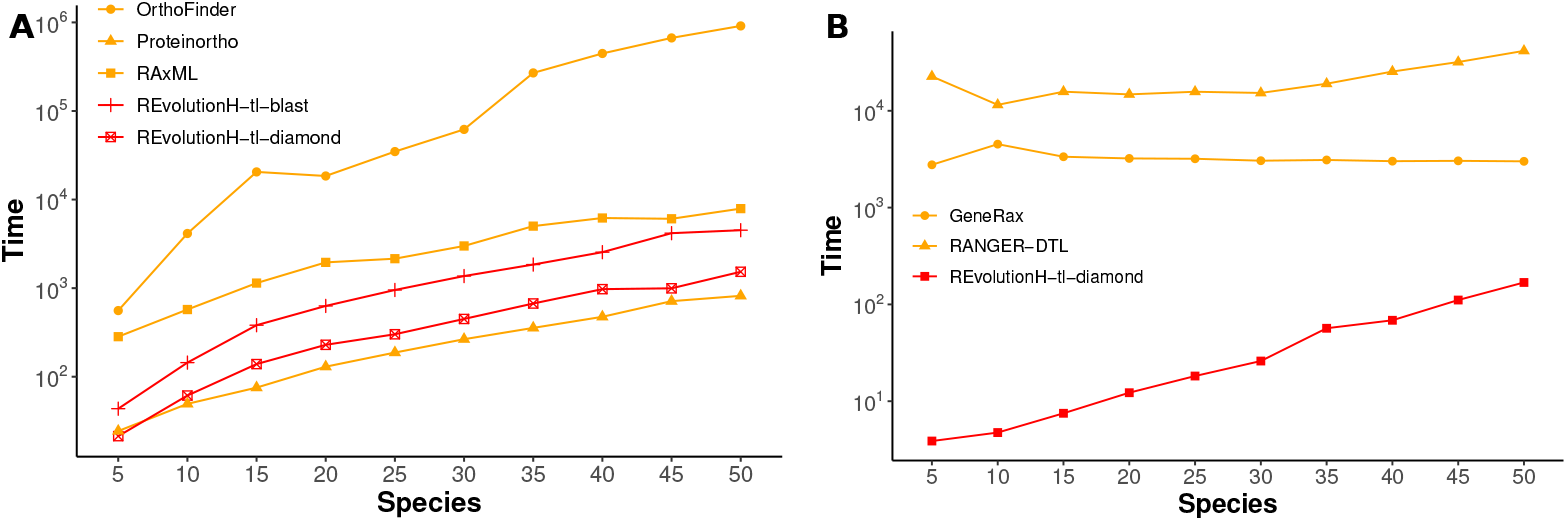
Running times. These figures show the total execution time needed for the analysis of the `SaGePhy5-50` dataset, measured in seconds. (A) Time performance for gene tree reconstruction and orthology inference. All the tools for this analysis were executed using 16 cpus. For `REvolutionH-tl`we only consider running times of steps one to three. (B) Time performance for three reconciliation. We consider only step 6 of `REvolutionH-tl`workflow.

In comparison with other tools, `REvolutionH-tl`demonstrates superior runtime performance relative to both `RAxML`and `Orthofinder`. As expected, `Proteinortho` completes its task more quickly than `REvolutionH-tl`; however, this speed comes at the cost of limited functionality, as `Proteinortho`only predicts orthology relations and does not infer gene or species phylogenies, nor perform reconciliations. Figure 10B presents the runtimes specifically for the reconciliation step. In this comparison, `REvolutionH-tl`clearly outperforms both `RANGER-DTL`and `GeneRax`, highlighting its efficiency in producing evolutionary scenarios while maintaining high accuracy.

## Discussion

`REvolutionH-tl` introduces a novel, efficient, and comprehensive framework for reconstructing evolutionary histories from sequence data. Unlike traditional methods that require precomputed gene or species trees and multiple specialized tools, `REvolutionH-tl` delivers an end-to-end pipeline that integrates orthology prediction, phylogenetic inference, and reconciliation into a unified system. Its core relies on the theoretical foundation of best match graphs (BMGs), allowing robust orthogroup and orthology detection through graph-based heuristics, which are both scalable and accurate.

Across benchmarking analyses, `REvolutionH-tl` consistently demonstrates high performance in reconstructing orthogroups and inferring orthology relationships. It matches or exceeds the accuracy of widely used tools such as `Orthofinder`and `Proteinortho`while maintaining a significantly reduced computational footprint.

Particularly in large simulated datasets with known ground truth, `REvolutionH-tl` recovers nearly all true orthogroups with minimal merging or splitting errors—an essential quality for downstream evolutionary analyses. The use of BMGs ensures that orthology is inferred based on evolutionary proximity rather than mere sequence similarity, mitigating common pitfalls in reciprocal best-hit approaches.

A central innovation of `REvolutionH-tl` lies in its inference of event-labeled gene trees and a de novo species tree, guided by orthology assignments rather than alignment likelihoods. Despite not using maximum likelihood or Bayesian methods, `REvolutionH-tl` produces gene trees with comparably low false-positive rates, as observed in evaluations against tools like `RAxML`and `GeneRax`. Furthermore, the tool’s reconciliation procedure consistently maps gene trees to species trees with high accuracy. Notably, reconciliation distances between inferred and true scenarios are lowest for `REvolutionH-tl`, affirming the biological plausibility of the reconstructed histories.

Beyond accuracy and efficiency, `REvolutionH-tl` sets itself apart as the first platform to provide built-in, publication-quality visualizations that facilitate the interpretation of evolutionary histories. Through intuitive plots, users can explore genome complexity, orthogroup statistics, and gene tree reconciliations in both global and orthogroup-specific contexts. These visualizations enhance the interpretability of results by embedding gene trees within species tree frameworks, clearly annotating events such as gene duplications, speciations, and losses. This level of graphical integration is unmatched by other tools, which often require manual scripting or external software to generate comparable insights.

The tool’s visual capabilities serve not only as a means of validation but also as a powerful communication aid, making it easier for biologists to trace evolutionary trajectories and gene family dynamics. With commands like plot summary and plot reconciliation, users can quickly generate diagrams that summarize gene content evolution, orthology types, and lineage-specific innovations—key features for comparative genomics and evolutionary studies.

In summary, `REvolutionH-tl` advances the field by delivering an integrated, graph-driven, and visually rich platform for evolutionary inference. Its balance of theoretical rigor, practical speed, and interpretability makes it a valuable resource for researchers exploring gene family evolution at scale. Future work will expand support for modeling horizontal gene transfer and enhance sensitivity to highly divergent sequences, further cementing `REvolutionH-tl`’s role as a cornerstone in computational evolutionary biology.

## Acknowledgments

Authors would like to thank Carlos A. González-Castro, Arlette España-Tinajero, Alejandro Flores-Lamas and Jesús Andrés Tinajero-Arteaga for technical support.

